# An Automated Pipeline for Differential Cell Counts on Whole-Slide Bone Marrow Aspirate Smears

**DOI:** 10.1101/2022.05.26.493480

**Authors:** Joshua E. Lewis, Conrad W. Shebelut, Bradley R. Drumheller, Xuebao Zhang, Nithya Shanmugam, Michel Attieh, Michael C. Horwath, Anurag Khanna, Geoffrey H. Smith, David A. Gutman, Ahmed Aljudi, Lee A.D. Cooper, David L. Jaye

**Affiliations:** Department of Pathology and Laboratory Medicine, Emory University, Atlanta, GA; Department of Biomedical Informatics, Emory University, Atlanta, GA; Department of Pathology, Northwestern University, Chicago, IL; Winship Cancer Institute, Emory University, Atlanta, GA

## Abstract

Pathologic diagnosis of bone marrow disorders relies in part on microscopic analysis of bone marrow aspirate (BMA) smears and manual counting of marrow nucleated cells to obtain a differential cell count (DCC). This manual process has significant limitations, including analysis of only a small subset of optimal slide areas and nucleated cells, and inter-observer variability due to differences in cell selection and classification. To address these shortcomings, we developed an automated machine learning-based pipeline for obtaining 11-component DCCs on whole-slide BMAs. This pipeline utilizes a sequential process of identifying optimal BMA regions with high proportions of marrow nucleated cells, detecting individual cells within these optimal areas, and classifying these cells into one of 11 DCC components. Convolutional neural network models were trained on 396,048 BMA region, 28,914 cell boundary, and 1,510,976 cell class images from manual annotations. The resulting automated pipeline produces 11-component DCCs that demonstrate high statistical and diagnostic concordance with manual DCCs among a heterogeneous group of testing BMA slides with varying pathologies and cellularities. Additionally, we show that automated analysis can reduce intra-slide variance in DCCs by analyzing the whole slide and marrow nucleated cells within optimal regions. Finally, pipeline outputs of region classification, cell detection, and cell classification can be visualized using whole-slide image analysis software. This study demonstrates the feasibility of a fully-automated pipeline for generating DCCs on scanned whole-slide BMA images, with the potential for improving the current standard of practice for utilizing BMA smears in the laboratory analysis of hematologic disorders.

## INTRODUCTION

The pathologic diagnosis of many benign and neoplastic hematologic disorders relies on examination of bone marrow aspirates (BMAs), liquid samples containing marrow nucleated cells^1,2^. Differential cell counts (DCCs) from microscopy on Wright-stained BMA smears are obtained by manually counting a set number of nucleated cells and recording the relative proportion of various cell types^3^. BMA DCCs yield insight into disease pathophysiology and can provide disease-defining information, notably for myeloid diseases (e.g., acute myeloid leukemia and myelodysplastic syndromes) and plasma cell neoplasms (e.g., multiple myeloma)^4-6^.

Current labor-intensive, manual analysis of BMA smears is confounded by many shortcomings which affect DCCs and potentially the final diagnosis. While a typical BMA may contain tens of thousands of marrow nucleated cells, only a subset of cells is counted, thereby increasing the statistically expected variability^7,8^. Additionally, given the use of such a small proportion of available cells, only a subset of optimal BMA slide regions is employed^3^. Since an aspirate sample is not perfectly homogeneous, consistent with functionally and biologically distinct marrow microniches, this subjectivity in localization for cell counting could impact measured relative cell proportions^9,10^. Finally, inter-observer variability in the cytologic classification of individual cells can lead to differences in DCCs^11,12^.

Previous studies have attempted to address these shortcomings by developing automated machine learning-based models for detection and classification of marrow nucleated cells from BMA smears. Wang et al. utilized a faster region-based convolutional neural network (Faster R-CNN) for detection of marrow nucleated cells in BMA smears^13^. We previously utilized Faster R-CNN and CNN networks to detect and classify cells in manually-annotated BMA regions, demonstrating strong performance on non-neoplastic BMAs^14^. Fu et al. compared automated classification results to manual DCCs and obtained strong correlation for three cell types^15^. Matek et al. and Yu et al. developed large expert-annotated training datasets to train highly accurate CNN-based cell classification models^16,17^. Nonetheless, most prior studies fall short of fully automating BMA analysis by failing to identify optimal BMA regions in an automated fashion, outputting full DCCs, and, importantly, directly comparing model outputs to clinically-reported manual DCCs.

Here, we present a fully-automated pipeline for obtaining 11-component DCCs from scanned whole-slide BMA smears (**Figure 1**). This pipeline consists of three sequential machine learning models which identify optimal regions on slides for cell counting, detect individual marrow nucleated cells within these regions, and subsequently classify these cells. The pipeline output includes a complete DCC of 11 cell types that can be directly compared to manual DCCs. This study demonstrates the strong performance of each of the three sequential steps in the pipeline, as well as the pipeline’s overall accuracy in yielding DCCs across heterogeneous samples of BMA smears from multiple different pathologies and with varying cellularities. Additionally, we show the capability of automated DCCs to reduce imprecision though whole slide analysis. Taken together, our results suggest a feasible automated pipeline for BMA DCCs with potential clinical utility in improving hematopathology diagnostics.

**Figure 1.**
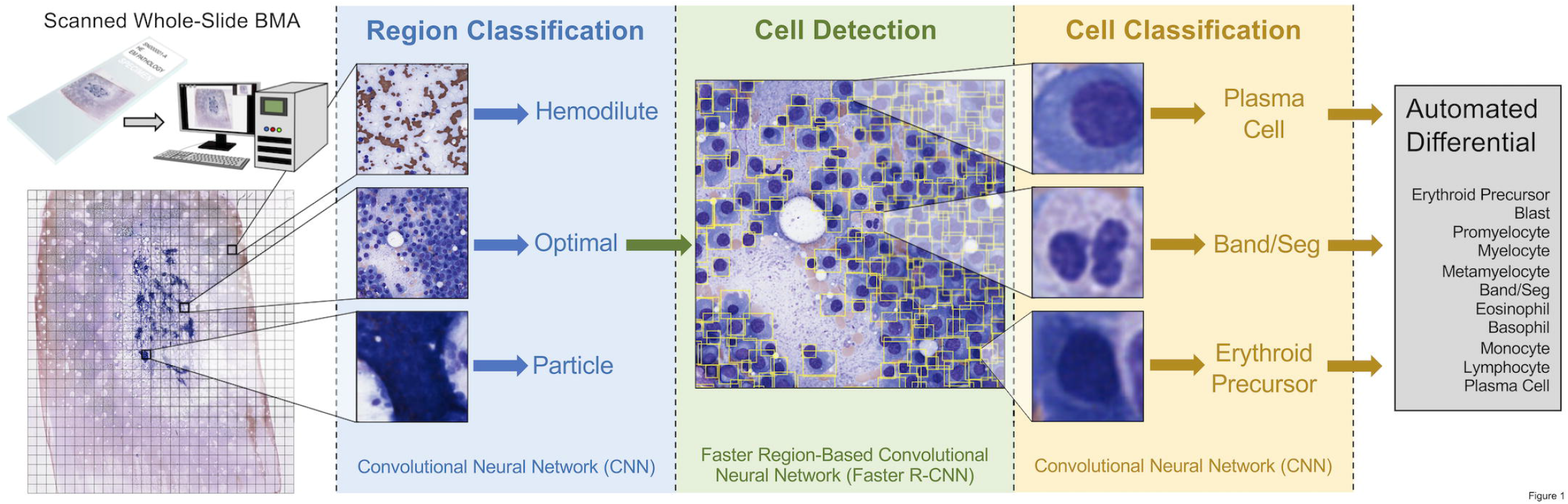
Overall workflow for automated machine learning-based pipeline for differential cell counts (DCCs) on whole-slide bone marrow aspirate (BMA) smears. BMA slides are scanned at 40x magnification (0.25 μm/pixel), and the scanned image is split into individual tiles. Each tile is processed by an EfficientNetV2S convolutional neural network (CNN) to determine whether it is an optimal tile to use for cell counting, or whether it represents a bone marrow particle, hemodilute region, or glass-only region outside of the aspirate. Each optimal tile is processed by a faster region-based convolutional neural network (Faster R-CNN) to detect individual cells within the tile, and each detected cell is processed by an EfficientNetV2L CNN to classify it as one of the cell differential components. All identified cells are averaged to output the final automated 11-component DCC.

## MATERIALS AND METHODS

### Bone marrow aspirate smears

Wright-stained BMA smears were uniformly prepared from EDTA-anticoagulated samples in the bone marrow laboratory at Emory University Hospital for routine patient care. Slides were scanned at 0.25 μm/pixel (40x) using a Leica Aperio AT2 scanner™. A spectrum of BMA smear slides with varying cellularities, pathologies, and white blood cell (WBC), red blood cell (RBC), and platelet counts, were chosen. **Supplementary Table 1** lists all BMA slides utilized in training and testing of machine learning models, and those used in prior work^14^. Whole-slide images (WSIs) were uploaded to a Digital Slide Archive server, and HistomicsUI was used for slide visualization and region/cell annotation^18^.

### Region classification

A training dataset for BMA region classification was generated from 69 BMA slides (**Supplementary Table 1**). On each WSI at 40x magnification, rectangular bounding boxes were manually drawn encompassing one of four region classes: 1) “optimal” – regions near aspirate particles with the highest proportions of marrow nucleated cells; 2) “particle” – regions containing dense aspirate particles; 3) “hemodilute” – bloody regions with high proportions of RBCs; and 4) “outside” – regions containing glass only. 10,948 regions of size at least 448×448 px were annotated (4,856 optimal, 2,644 particle, 3,318 hemodilute, 130 outside).

Each annotated region was cropped into multiple 448×448 px images, where the number of cropped images from each region was determined as follows:

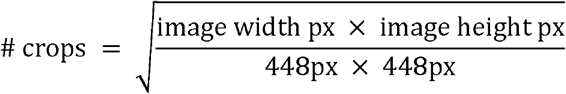

Each cropped image was copied 16 times, and each copy randomly augmented by flipping horizontally and/or vertically, rotating by a random multiple of 90°, and randomly changing brightness (0.75x-1.25x original) and contrast (0.9x-1.4x original). In total, 396,048 cropped/augmented region images were obtained (119,456 optimal, 58,528 particle, 176,672 hemodilute, 41,392 outside).

A sequential neural network architecture consisting of an EfficientNetV2S backbone CNN with weights pre-trained on ImageNet, global average pooling layer, batch normalization layer, dropout layer (dropout rate = 0.2), and dense layer with softmax activation function, was used to classify regions into one of four classes^19^. First, all layers except the top layers were frozen. the top layers were trained with a larger learning rate (Adam optimizer with learning rate = 1e-3) and categorical cross-entropy loss function. Subsequently, all layers were unfrozen, and the entire model was trained using a smaller learning rate (1e-5). Ten-fold cross-validation was performed, with data split into training (64%), early stopping (16%; patience of 10 epochs), and validation (20%) datasets. For each cross-validation split, the area under the receiver operating characteristic (AUROC) performance metric was calculated.

### Cell detection

A training dataset for BMA cell detection was generated from 36 BMA slides (**Supplementary Table 1**). BMA slide regions of varying cellularities containing cells with distinct cell boundaries were identified. Rectangular bounding boxes were drawn around all cells in these areas at 40x magnification. All cell types used for cell classification were included. In total, 28,914 cell bounding boxes within 215 BMA areas were obtained.

HistomicsDetect was used to develop a faster region-based convolutional neural network (Faster R-CNN) for cell detection^20,21^. This network utilized a ResNet50V2 CNN backbone with 14 residual blocks^22^. Tau (minimum intersection over union (IoU) to validate a detection) and nms_iou (maximum IoU allowed between detections) parameters were set to 0.5 and 0.1, respectively.

HistomicsDetect provides significant advantages over prior cell detection packages, including utilizing ROIAlign versus ROI pooling, improving selection of anchor sizes to match cell sizes and shapes, and decoupling of cell detection and classification with two separate models, allowing for collection of Supplementary annotations for rarer cell types outside of detection ROIs for more efficient cell classifier training^21,23^. Ten-fold cross-validation was performed, with data split into training (64%), early stopping (16%; patience of 10 epochs), and validation (20%) datasets. For each cross-validation split, average precision with IoU thresholds of 0.25, 0.50, and 0.75 (AP25, AP50, and AP75, respectively) performance metrics were calculated.

### Cell classification

A training dataset for BMA cell classification was generated from 73 BMA slides (**Supplementary Table 1**). Points representing cell centroids were annotated at 40x magnification, and cell class was recorded from one of 16 labels: erythroid precursor, blast, promyelocyte, myelocyte, metamyelocyte, band/seg, eosinophil, basophil, monocyte, lymphocyte, plasma cell, macrophage, mast cell, megakaryocyte, unknown intact cell, and disrupted cell. Neoplastic and non-neoplastic counterparts (e.g., neoplastic and benign blasts) were grouped as one class. 23,609 cells were annotated, and cell images were extracted by drawing 64×64 px bounding boxes around cell centroids. Each cell image was copied 64 times, with each copy undergoing random center cropping (central fraction between 0.7-1.0) and resizing to 64×64px, horizontal and/or vertical flipping, rotation by a random multiple of 90°, and random changes to image brightness (0.75x-1.25x original) and contrast (0.9x-1.4x original). In total, 1,510,976 augmented cell images were produced.

A sequential neural network architecture consisting of an EfficientNetV2L backbone CNN with weights pre-trained on ImageNet, global average pooling layer, batch normalization layer, dropout layer (dropout rate = 0.2), and dense layer with softmax activation function, was used to classify cell images into one of sixteen classes^19^. Model training and validation were performed analogously to region classification.

### WSI BMA pipeline

We employed 44 BMA slides not used in the training of region classification, cell detection, or cell classification models for testing the automated pipeline (**Supplementary Table 1**). For all testing slides, a manual 11-component DCC (300 or 500 cells total) had been performed per ICSH guidelines by a medical technologist or hematopathologist using glass slides ^3^. Each BMA WSI was split into 448×448 px tiles. The predicted region class of each tile was determined by the region classification model. Optimal tiles were then subjected to cell detection, and identified cells were processed by the cell classification model. Cells with a predicted class within the 11-component DCC (erythroid precursor, blast, promyelocyte, myelocyte, metamyelocyte, band/seg, eosinophil, basophil, monocyte, lymphocyte, and plasma cell) underwent further analysis, whereas other detected cells (macrophage, mast cell, megakaryocyte, unknown intact cell, and disrupted cell) did not. Automated DCC percentages were calculated by dividing the number of identified cells of a particular cell type by the total number of identified cells among all cell types within the 11-component DCC. Test-time augmentation was performed at region and cell classification steps, with 16 and 64 random augments per image being utilized, respectively.

When comparing manual and automated DCC percentages among testing slides, the Pearson correlation coefficient (*r*) and concordance correlation coefficient (ρ_*c*_) were calculated:

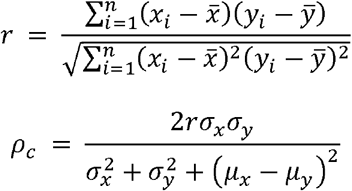

Statistically expected variability in manual and automated DCC percentages was determined by calculating 95% confidence intervals based on the binomial distribution^8^.

### Localized differentials

Localized DCCs were calculated at the location of each detected marrow nucleated cell on the BMA WSI with a predicted class within the 11-component DCC. For each starting cell, the 499 nearest neighboring cells that had a predicted class within the 11-component DCC were identified. A 500-cell DCC was then calculated using the cell classes of the starting cell and these 499 nearest neighbors. For visualization of localized DCCs, differential values were averaged across all starting cells located within an individual tile.

### Software and hardware

TensorFlow 2.8.0 was used to develop and train neural network models^24^. All experiments were run on a Linux server with Intel Xeon Gold 6132 processor, 126 GB RAM, and 4 Tesla V100-PCIE-32GB GPUs.

## RESULTS

### Region classifier accurately identifies optimal areas for DCCs

Previous studies relied on manual identification of optimal BMA slide areas for further cell detection and classification^14,15^. Here, we developed a machine learning model for automating optimal region identification. This model classifies BMA regions as one of four types: 1) “optimal” – regions richest in marrow nucleated cells, ideal for the DCC, and also called cellular trails; 2) “particle” – regions containing dense aspirate particles; 3) “hemodilute” – regions of mostly blood; and 4) “outside” – glass-only regions. To develop a training dataset for BMA region classification, rectangular bounding boxes encompassing these four region types were manually annotated among 69 designated training BMA slides (**Figure 2A; Supplementary Table 1**). To increase training data sample size and model performance, image cropping and augmentation (flip, rotation, brightness/contrast adjustment) was performed on each annotated region. In total, 10,948 regions were annotated, resulting in a training dataset of 396,048 total cropped/augmented images for model training (**Figure 2B**).

**Figure 2.**
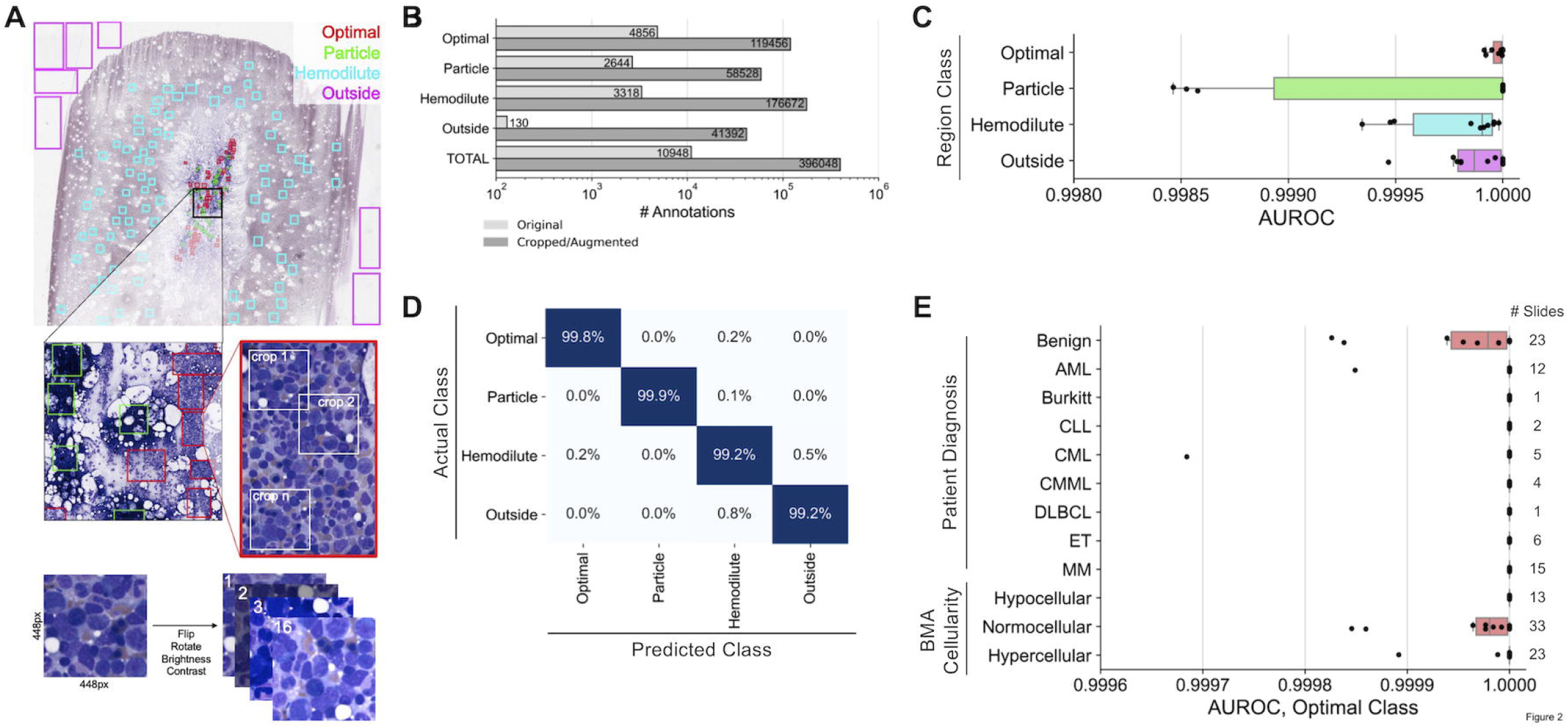
Training and evaluation of the bone marrow aspirate region classifier. **(A)** Generation of training data for the region classifier. Within each training slide, rectangular regions are manually drawn consisting of one of four classes (optimal, particle, hemodilute, outside). Random 448×448 pixel crops of each region are made, and each crop is subjugated to 16 iterations of augmentation, including flipping, rotation, brightness adjustment, and contrast adjustment. **(B)** Number of annotated regions before and after region cropping/annotation for model training. **(C)** Performance of the region classifier across 10 cross-validation splits. Each point represents a cross-validation split. **(D)** Confusion matrix of predictions from the region classifier cross-validation. **(E)** Region classifier performance within slide subsets representing different patient diagnoses and aspirate cellularities. Each point represents a cross-validation split. AUROC: area under the receiver operating characteristic.

Overall, the region classification model demonstrated strong performance on all four region classes, with average AUROC values across ten cross-validation folds exceeding 0.9995 for each class (**Figure 2C**). The optimal class showed particularly strong performance, with a mean AUROC of 0.99997, mean sensitivity of 99.79%, and mean specificity of 99.89%. Misclassifications were minimal, with the largest misclassifications being between hemodilute and outside classes, and only 22.9% of misclassifications involving optimal tiles (**Figure 2D**). Additionally, performance was excellent on slides across different patient diagnoses, including those with high WBC and platelet counts, and marrow cellularities, with mean optimal class AUROC values above 0.9999 for all subsets (**Figure 2E**). These results demonstrate that the machine learning-based region classification model accurately identifies optimal BMA slide regions for further detection and classification of marrow nucleated cells without regard to overall smear cellularity, blood composition, or pathologic diagnosis.

### Cell detector and classifier accurately identify marrow nucleated cells

To develop a training dataset for BMA cell detection, 215 areas among 36 designated training BMA slides with varying cellularities and patient pathologies were manually identified; within these 215 areas, bounding boxes around 28,914 identified cells were manually drawn (**Figure 3A; Supplementary Table 1**). This training dataset represents a significant improvement over previous BMA cell detection studies by including an order of magnitude larger number of cell boundary annotations as well as incorporating non-neoplastic and neoplastic cases^13,14^. The cell detection model demonstrates strong performance as evidenced by average precision-at-intersection-over-union (IoU) threshold of 0.5 (AP50) values matching or exceeding those in similar cell detection studies (**Table 1**)^13,25^. AP50 values did not substantially differ between samples representing different pathologic diagnoses or between slides with varying cellularities (**Table 2**).

**Table 1.** 10-fold cross-validation metrics for the cell detection model.

| **Metric** | **AP25** | **AP50** | **AP75** |
| --- | --- | --- | --- |
| **Mean** | 0.930 | 0.890 | 0.672 |
| **Standard Error** | 0.003 | 0.003 | 0.009 |

2 *AP25:* Average precision with intersection over union (IoU) threshold of 0.25
3 *AP50:* Average precision with intersection over union (IoU) threshold of 0.50
4 *AP75:* Average precision with intersection over union (IoU) threshold of 0.75

**Table 2.** Cell detection AP50 metrics for BMA slide subsets.

| **Slide Subset** | **AP50, Mean** | **AP50, Standard Error** |
| --- | --- | --- |
| Diagnosis – Benign | 0.871 | 0.005 |
| Diagnosis – AML | 0.923 | 0.006 |
| Diagnosis – MM | 0.903 | 0.005 |
| Cellularity – Hypocellular | 0.914 | 0.006 |
| Cellularity – Normocellular | 0.878 | 0.004 |
| Cellularity – Hypercellular | 0.902 | 0.006 |

**Figure 3.**
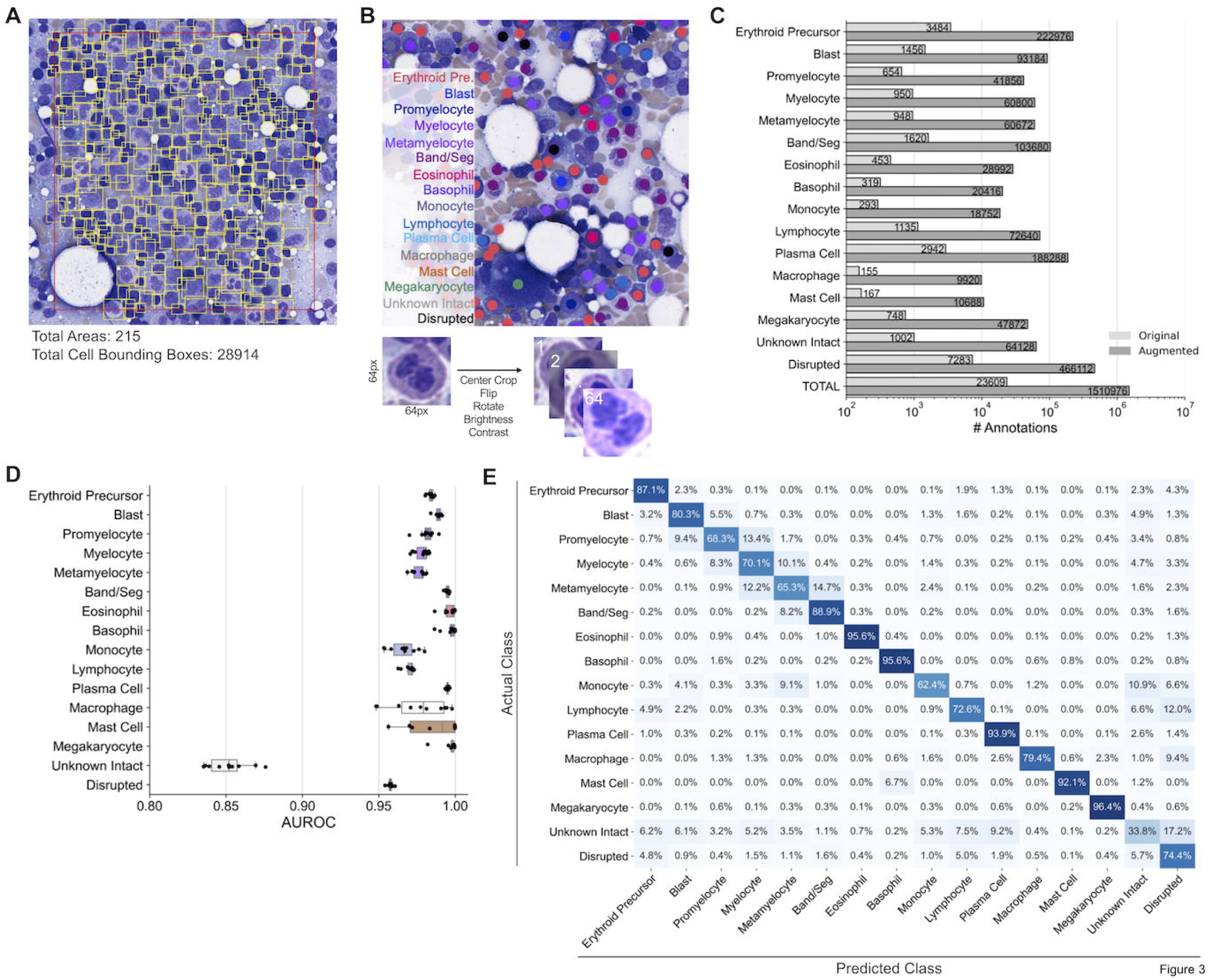
Training and evaluation of the bone marrow aspirate cell detector and classifier. **(A)** Generation of training data for the cell detector. Within each training slide, optimal areas are chosen and bounding boxes around each cell within these optimal areas are manually drawn. The number of annotated areas and cell bounding boxes for model training are provided. **(B)** Generation of training data for the cell classifier. Within each training slide, points are manually drawn at the centroid of cells, with the cell type of each manually-classified cell being noted. For each classified cell, a 64×64 pixel bounding box is made around the centroid of each cell, and each cell is subjected to 64 augmentations, including central cropping, flipping, rotation, brightness adjustment, and contrast adjustment. **(C)** Number of annotated cell classifications before and after image augmentation for model training. **(D)** Performance of the cell classifier across 10 cross-validation splits. Each point represents a cross-validation split. **(E)** Confusion matrix of predictions from the cell classifier cross-validation. AUROC: area under the receiver operating characteristic.

To develop a training dataset for BMA cell classification, 23,609 cells among 16 cell classes were manually annotated from 73 designated training BMA slides (**Figure 3B; Supplementary Table 1**). Cell images centered at labeled cell centroids underwent further augmentation (center crop, flip, rotation, brightness/contrast adjustment) to increase training data sample size and model performance. In total, 1,510,976 total augmented images were utilized for model training (**Figure 3C**). The cell classification model demonstrated strong performance for all 11 cell types within the DCC, and 3 additional cell types (macrophage, mast cell, megakaryocyte), evidenced by mean AUROC values exceeding 0.95 (**Figure 3D**). While the model also performed well on disrupted cells, lower performance was observed for unknown intact cells, likely because this class represents a heterogeneous catch-all group for intact cells with ambiguous morphology. The model most accurately identified cells with very distinct morphologies, including eosinophils, basophils, plasma cells, and megakaryocytes (**Figure 3E**). As expected, some misclassifications were made stepwise along the myeloid lineage, for example between blasts and promyelocytes, promyelocytes and myelocytes, etc. While the overall accuracy of unknown intact cell classifications was poor, the relatively uniform misclassification of these cells across all other classes within the 11-component DCC indicates that these misclassifications are unlikely to significantly impact automated differential outputs. Overall, this cell classification model matches the performance of prior BMA cell classification studies which either only utilized cells from non-neoplastic BMA slides or required significantly greater annotations for model training^14,16,17^.

### Automated pipeline outputs DCCs with high correspondence to manual DCCs

Trained region classification, cell detection, and cell classification models were combined to develop an automated sequential pipeline which outputs an 11-component DCC from a BMA whole slide image. Outputted automated differentials from the pipeline were compared to manual DCCs using 44 testing BMA slides that were not utilized in model training. An average of 19,753 viable cells, excluding “unknown intact” and “disrupted” classes, were detected and classified from optimal regions, with a range of 237 to 126,483 cells per slide (**Figure 4A**). **Figure 4B** shows the regression of erythroid precursor differential percentages between manual and automated DCCs among testing slides. **Supplementary Figure 1** displays regressions for all 11 cell types. Most cell types had strong correlation based on the Pearson correlation coefficient *r* (measure of linearity) and the concordance correlation coefficient ρ_*c*_ (measure of regression along the 1:1 line); correlation coefficients were similarly strong when combining all cell classes on single slides (**Figure 4C**). Better performance was observed for more common cell types (erythroid precursors, band/segs, lymphocytes) and those with significant diagnostic relevance (blasts, plasma cells), with lower performance on myeloid lineage maturation intermediates (myelocytes, metamyelocytes) and rarer cell types (basophils, monocytes). Bland-Altman plots show minimal bias for most cell types, with exceptions including blasts and plasma cells, which tended to be underestimated by automated DCCs for AML and MM cases containing high proportions of these cell types, respectively (**Supplementary Figure 2**). We found that these biases were due to a proportion of neoplastic blasts and plasma cells in these cases being classified as “unknown intact” cells and thus not being incorporated into the 11-component DCC (**Figure 3E; Supplementary Figure 3**); however, biases appear to be largest in cases for which both manual and automated DCC values still exceed clinically relevant thresholds and thus would not impact pathologic diagnosis.

**Figure 4.**
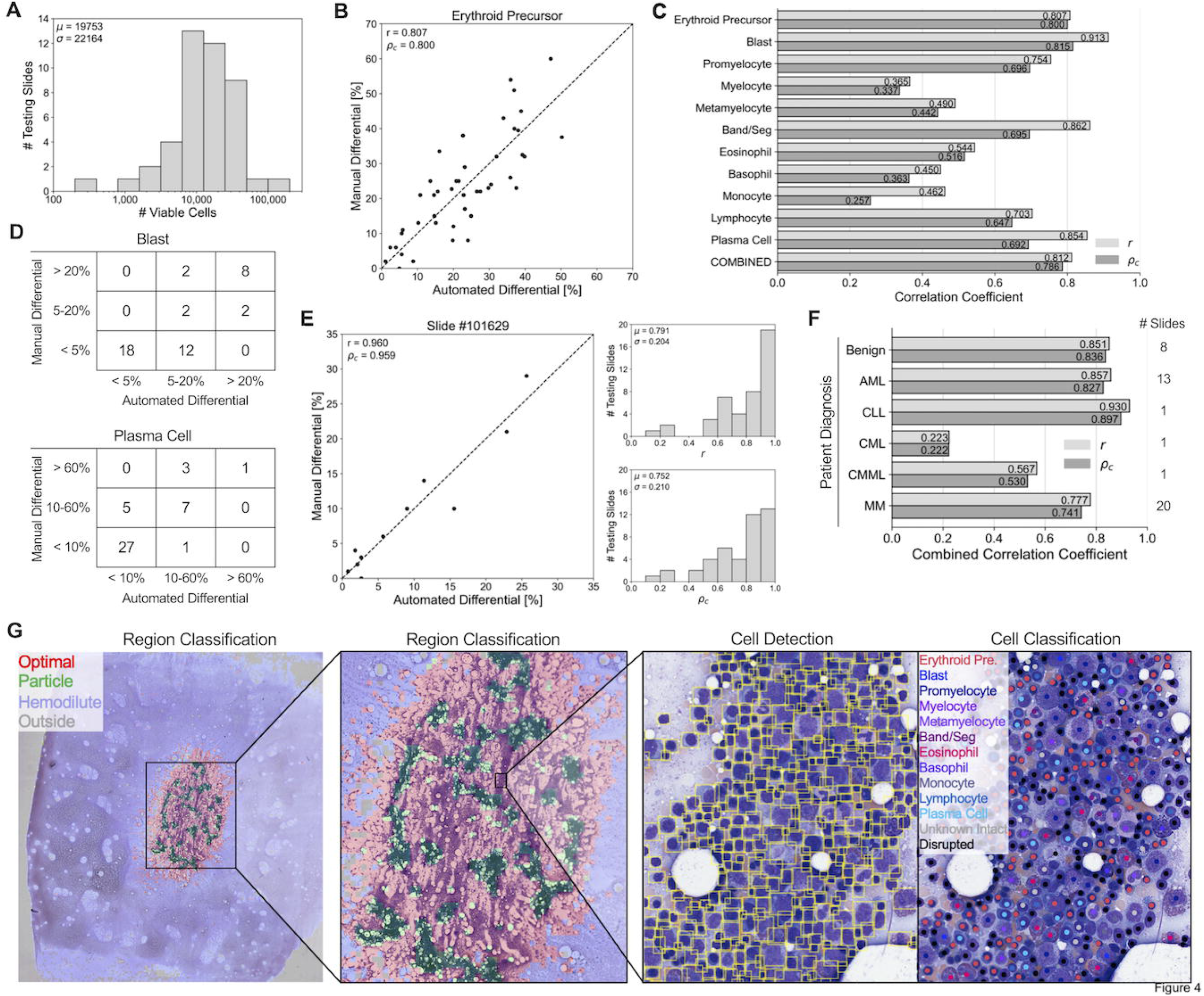
Evaluation of the automated machine learning-based pipeline for DCCs on whole-slide bone marrow aspirate smears. **(A)** Histogram showing the number of viable cells identified by the automated pipeline for each of 44 testing slides. **(B)** Correlation plot comparing the percentage of erythroid precursors obtained from the manual differential, versus the percentage obtained from the automated machine learning-based pipeline averaged across all identified viable cells in each slide. Each point represents one of 44 testing slides. Dotted line represents a 1:1 correlation between manual and automated values. *r*: Pearson correlation coefficient between manual and automated values. _/li*c*_: concordance correlation coefficient between manual and automated values, representing how well points are fitted by the 1:1 correlation line. **(C)** Pearson and concordance correlation coefficients for each cell type, as well as the combined correlation coefficients across all cell types. **(D)** 3×3 contingency tables for the number of slides with manual and automated differential percentages of (Top) blast cells below 5%, between 5-20% and above 20%; and (Bottom) plasma cells below 10%, between 10-60%, and above 60%. **(E)** (Left) Correlation plot comparing the percentage of 11 different cell types from the manual 11-component DCC of an individual slide, versus the percentages obtained from the automated machine learning-based pipeline. Each point represents one of 11 cell types. Dotted line represents a 1:1 correlation between manual and automated values. (Right, Top) Histogram showing the individual slide-based Pearson correlation coefficients for each of 44 testing slides. (Right, Bottom) Histogram showing the individual slide-based concordance correlation coefficients for each of 44 testing slides. **(F)** Pipeline performance within slide subsets representing different patient diagnoses. **(G)** Visualization of pipeline outputs for an individual testing slide. (Left, Middle) Region classification outputs. (Right) Cell detection and classification outputs.

We next compared results of automated and manual DCCs using clinically employed thresholds for myeloid neoplasms (5% and 20% blasts) and plasma cell neoplasms (10% and 60% plasma cells) (**Figure 4D**)^1^. While few discrepancies between automated and manual DCCs were identified with blast percentages at the 20% threshold (40/44, 90.9% agreement), a larger proportion of discrepancies were found at the 5% threshold (32/44, 72.7% agreement). Similarly, fewer discrepancies in plasma cell percentages were observed at the 60% threshold (41/44, 93.2%) compared to the 10% threshold (38/44, 86.4% agreement).

Strong correlation between manual and automated DCCs for individual BMA slides was observed when regressing across all 11 cell types (**Figure 4E; Supplementary Figure 4**). BMA slides from benign, AML, and CLL cases demonstrated the highest correlation, with relatively poor correlation on the two CML cases analyzed (**Figure 4F**). **Figure 4G** shows the visualization of pipeline outputs on an example WSI BMA slide. Overall, the automated machine learning-based pipeline provides 11-component DCCs with high correspondence to manual DCCs.

### Intra-slide and inter-observer variance confound manual DCC comparisons

Variance in manual BMA DCC values for a particular slide is expected between hematopathologists or even between multiple DCCs from the same hematopathologist; this variance has multiple sources, including counting only a small subset of cells on the slide (typically 500), utilization of only a subset of optimal BMA slide regions, and differences in cell classification between observers. Statistically expected variance in manual BMA DCC values as a function of number of counted cells has been previously assessed by calculating 95% confidence intervals based on the binomial distribution^8^. This variance is significantly smaller in automated DCC values compared to manual DCCs due to the much larger number of counted cells. Additionally, the large variance observed in manual DCCs greatly affects the comparison of manual and automated DCC values, both at the cell type level and slide level (**Figures 5A and 5B, Supplementary Figures 5 and 6**). If slides with variability large enough to overlap with clinically-relevant thresholds are excluded from analysis, greater agreement between manual and automated DCCs is observed at 3/4 threshold values analyzed in **Figure 4D** (**Figure 5C**; 20% blasts: 33/33, 100%; 5% blasts: 22/33, 66.7%; 60% plasma cells: 34/36, 94.4%; 10% plasma cells: 34/36, 94.4%).

**Figure 5.**
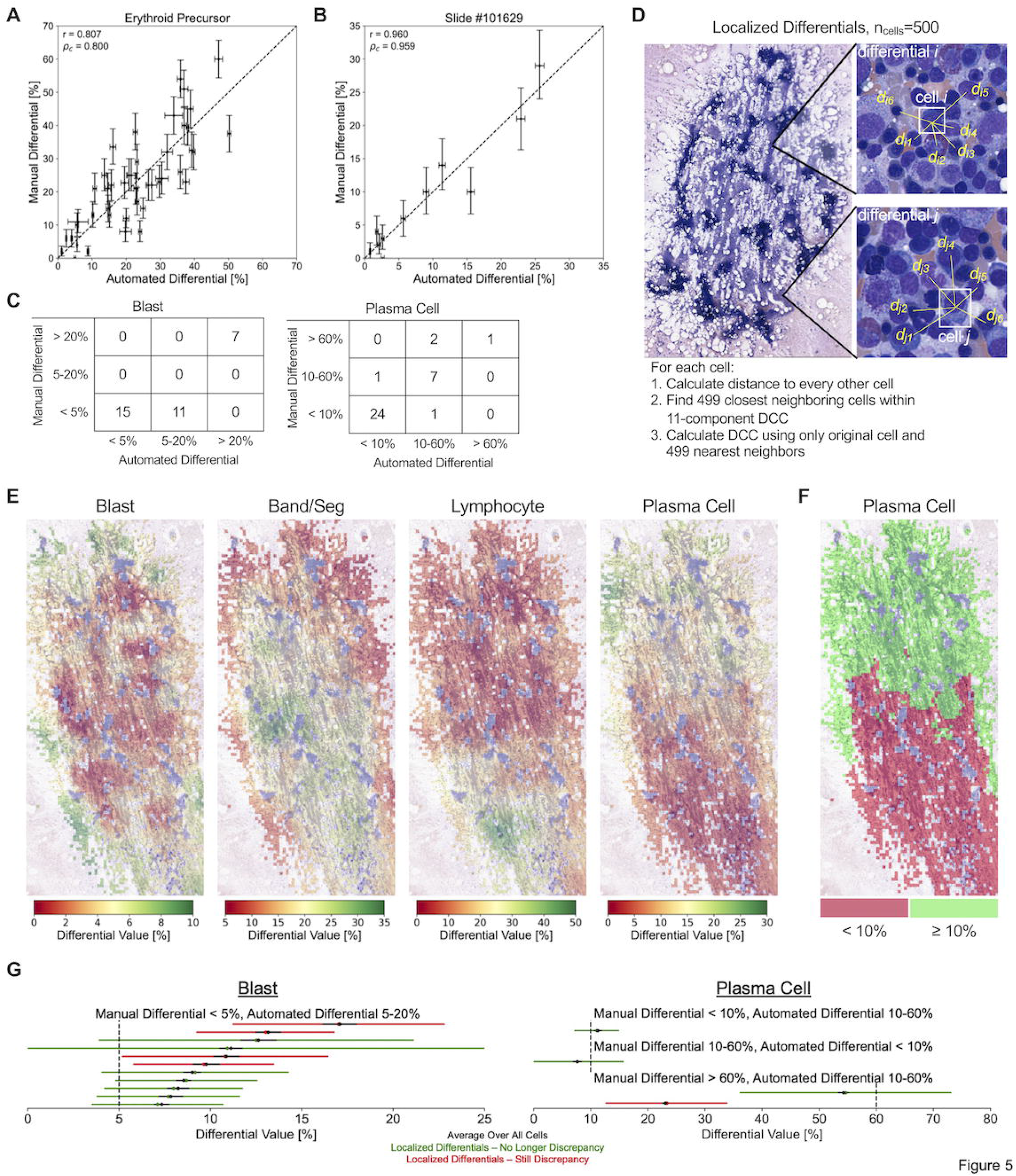
Assessment of intra-slide variance in DCC values. **(A)** Correlation plot comparing erythroid precursor differential values between manual and automated DCCs, with vertical and horizontal error bars represent the statistically expected variability in manual and automated differential values, respectively, based on the number of cells identified from the manual and automated methods. Each point represents one of 44 testing slides. Dotted line represents a 1:1 correlation between manual and automated values. *r*: Pearson correlation coefficient between manual and automated values. _/li*c*_: concordance correlation coefficient between manual and automated values, representing how well points are fitted by the 1:1 correlation line. **(B)** Correlation plot comparing the percentage of 11 different cell types between manual and automated DCCs, with vertical and horizontal error bars represent the statistically expected variability in manual and automated differential values, respectively, based on the number of cells identified from the manual and automated methods. Each point represents one of 44 testing slides. Dotted line represents a 1:1 correlation between manual and automated values. **(C)** 3×3 contingency tables for the number of slides with manual and automated DCC values below and above clinically-relevant thresholds, excluding slides with statistically expected variance overlapping with threshold values. **(D)** Diagram demonstrating the calculation of 500-cell localized differentials using each individual cell and their 499 nearest neighboring cells. **(E)** Visualization of the localized variability in DCC values across different optimal regions of an example BMA smear. For each cell, a localized 500-cell differential is calculated among the cell and its 499 nearest neighbors, and differential values are averaged across all cells located within an individual optimal tile. Optimal tiles are colored based on average tile values for a particular cell type. **(F)** Thresholding of localized DCC plasma cell values at the 10% diagnostic threshold for plasma cell neoplasms. **(G)** Intra-slide variability in localized differential values for (left) blast cells and (right) plasma cells among the 15 BMA slides with discrepancies between manual and automated values identified in **Figure 5C**. Each BMA slide is represented by a horizontal line. Black dot and line represent the mean value and statistically expected variability, respectively, in automated differential values when averaging across all cells. Green/red dot and line represent the mean value and 95% confidence interval, respectively, in localized differential values. BMA slides that no longer demonstrate a discrepancy between manual and automated values across the diagnostic threshold when taking into account intra-slide variability are shown in green; those that still demonstrate discrepancies are shown in red.

To further quantify intra-slide variance in DCC values and its impact on the comparison between manual and automated DCCs, we calculated 500-cell localized differentials using our computational pipeline (**Figure 5D**). These localized differentials assess local slide-level changes in DCC values by calculating a 500-cell DCC using individual cells and their 499 nearest neighboring cells; localized differentials were calculated at the location of each identified cell on the BMA slide, and the range in differential values across the entire slide was determined. **Figure 5E** provides a visualization of localized differential values, showing that peaks and troughs in DCC values are highly localized to particular slide areas. This localized variation can straddle clinically-relevant thresholds, including plasma cell levels above and below 10% (**Figure 5F**)^1^. We therefore used this tool as another method of measuring imprecision of the automated blast and plasma cell percentages. Notably, 10/15 testing slides presented in **Figure 5C** no longer displayed overt discrepancies between manual and automated DCCs at diagnostic thresholds when accounting for such intra-slide variance (**Figure 5G**). Furthermore, we also measured imprecision in the manual DCCs of the remaining 5 discrepant cases by assessing interobserver variability. We found inter-observer variations in the DCCs of 5 pathologists that crossed diagnostic thresholds for 3/5 cases, rendering these apparent discrepancies of less significance (**Supplementary Table 2**). These latter results also highlight intra-slide variance in manual DCC values which could have diagnostic ramifications.

### Automated pipeline minimizes intra-slide variance in DCC values

We hypothesized that our automated pipeline for DCCs provides two distinct advantages in reducing intra-slide variance for DCC values: (1) the pipeline assesses cells from all optimal BMA regions, whereas manual DCCs focus on small subsets of optimal regions; and (2) the pipeline analyzes all detected marrow nucleated cells, whereas manual DCCs analyze a significantly smaller set number of cells per slide, generally 500 or fewer. To quantify these advantages, we compared the variance in automated DCC values between 500-cell DCCs from localized BMA areas, and 500-cell and 5000-cell DCCs with cells taken randomly from all BMA regions (**Figure 6**). For all 11 cell types, statistical imprecision significantly decreased by switching from localized BMA areas to randomly-localized cells across the BMA slide, and by increasing analyzed cells from 500 to 5,000. This suggests that the location-agnostic approach and increased cell utilization from automated DCCs can substantially reduce intra-slide variance in DCC values observed from manual DCCs, and thus could lead to more representative assessments.

**Figure 6.**
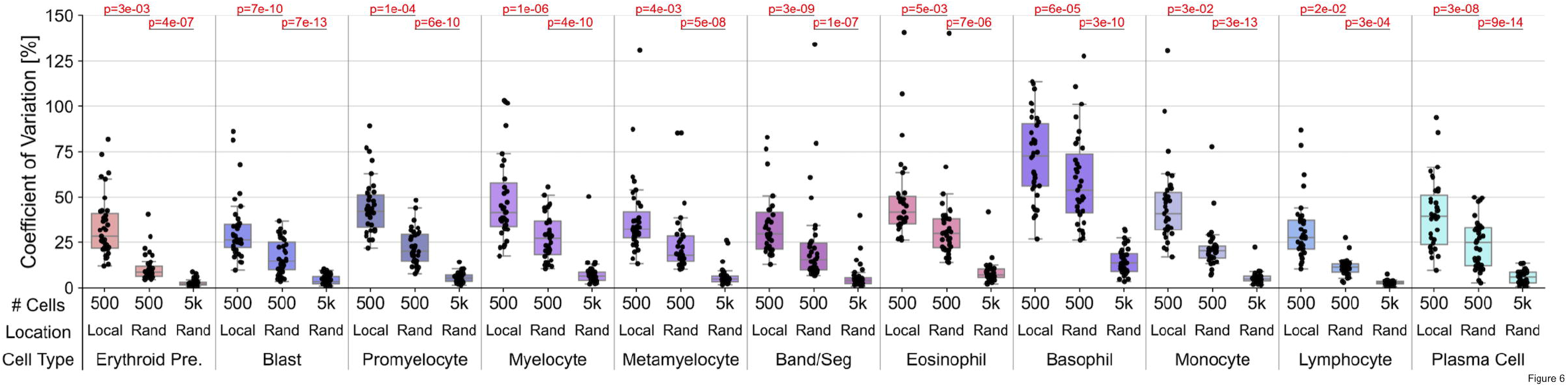
Reduction of intra-slide variance by automated DCCs. Comparison of coefficient of variation (standard deviation divided by mean) in BMA slide DCC values between 500-cell DCCs in localized (“Local”) slide areas, 500-cell DCCs with cells taken from random (“Rand”) slide areas, and 5000-cell DCCs with cells taken from random slide areas. P-values for two-tailed t-tests with unequal variance are provided.

## DISCUSSION

Most prior studies explored automation of BMA slide analysis by developing machine learning-based models for cell detection and classification^13-17^. However, these studies fall short of providing fully automated BMA DCCs by relying on manual selection of optimal regions for subsequent cell detection and classification, focusing on select portions of the BMA analysis pipeline, or restricting their application to non-neoplastic samples. Wang et al. incorporated a less selective region classifier in their model, but importantly did not compare outputs to manual DCCs^26^. Here, we present a fully-automated pipeline for DCCs from whole-slide BMA smears for benign and malignant samples (**Figure 1**). The pipeline encompasses a sequential process of classifying BMA slide regions, detecting marrow nucleated cells within optimal regions, and classifying these cells as one of 11 constituents of the DCC. This methodology employs a novel approach of automating the classification of slide regions into optimal, particle, hemodilute, or outside categories using a CNN model (**Figure 2**). The region classification model accurately identifies optimal regions across all subsets of BMA slides, including benign and a spectrum of malignant cases and of smear cellularities. Additionally, the cell detection and classification models demonstrated strong performance across all 11 cell types of the DCC and 3 other cell types (macrophages, mast cells, and megakaryocytes) (**Figure 3**). Of note, the performance of this cell classification model matches that reported in prior studies, which either restricted their dataset to non-neoplastic cells or utilized significantly more cell annotations, and is likely due to our implementation of extensive image augmentation^14,16,17,27,28^. Our sequential approach closely resembles the standard procedure for manual microscopic analysis, namely starting with a low-power view to identify optimal regions, followed by high-power cell assessment within these regions. Because of this close resemblance, we expect that this automated region classification approach could be applied to other areas of digital pathology^29-31^.

Automated DCCs from our integrated pipeline showed a promising degree of correspondence with manual DCCs at multiple levels: 1) regression of manual versus automated DCC percentages for individual cell types across all testing slides; 2) regression of DCC percentages for individual testing slides across all cell types; and 3) contingency tables of DCC percentages below versus above clinically-relevant thresholds when accounting for imprecision (**Figure 4**). Both cell-type performance and slide performance were highly correlated with training dataset composition, with greater correspondence observed for more common cell types (erythroid precursors, blasts, band/segs, plasma cells) and for diagnoses with greater representation in training datasets (benign, AML, MM). This suggests that pipeline performance can be directly improved by training dataset enlargement and diversification. Minimal bias in automated DCC percentages was observed for most cases, except for underestimation of blast and plasma cell percentages in AML and MM cases containing high proportions of these cell types, respectively (**Supplementary Figure 2**). These biases were due to a proportion of neoplastic blasts and plasma cells being misclassified as “unknown intact” cells by the automated pipeline, analogous to the clinical situation of how neoplastic cells with ambiguous morphology are often placed in an “other” category until ancillary studies are provided to assess neoplastic cell lineage and correctly re-categorize these cells (**Supplementary Figure 3**). Such misclassifications by the automated pipeline could potentially be improved upon by increasing the number of myeloid malignancy and MM cases in training datasets and by utilizing separate cell classes for non-neoplastic and neoplastic counterparts (e.g., non-neoplastic and neoplastic blasts).

A novel aspect of this work, in comparison with other reported machine learning approaches to BMA DCC analysis, is the introduction of slide-level method comparisons of automated outputs to clinically-obtained manual results and their imprecisions^15,26^. While the utility remains for traditional machine learning performance metrics, such as those shown in **Figures 2 and 3**, ultimate clinical laboratory validation of these automated tools will undoubtedly require extensive slide- and disease-level comparisons. In this promising study, strong correspondence between manual and automated DCC percentages was observed at 3 of 4 clinically-relevant thresholds for blasts and plasma cells (at least 86.4% agreement; **Figure 4D**). This correspondence was improved when taking into account statistically expected variability in manual DCC values (3/4 thresholds with at least 94.4% agreement; **Figure 5C**). Additionally, among the 15 slides which showed discrepancies between manual and automated DCC percentages at these thresholds, 10/15 no longer displayed discrepancies when accounting for intra-slide variance in localized differential values (**Figure 5G**), and 3/5 slides with continued discrepancies had significant inter-observer variations in manual DCCs that crossed diagnostic thresholds (**Supplementary Table 2**). Thus, while manual 500-cell DCCs remain the gold standard of analyzing BMA nucleated cells to which comparison of automated pipeline results should be made, intrinsic imprecision in manual DCCs should be considered in such method comparisons.

Future efforts will allow for enhanced generalizability and performance of machine learning-based models for automated BMA DCCs. Diverse BMA image datasets from multiple institutions with varying staining protocols and patient populations would ideally be combined to train machine learning-based models with strong multi-institution performance. Matek et al. developed a large publicly-available dataset of expert-annotated BMA cell images for training of CNN-based cell classification models^16,32^. Interestingly, but perhaps unsurprisingly, global differences in cell images were identified between their cell classification dataset and ours via unsupervised learning (**Supplementary Figure 7**), likely due in part to differences in staining protocols (Wright vs Pappenheim), and patient diagnoses (greater proportion of benign and AML cases vs MM cases) in training sets. Domain adaptation could be explored as a potential approach to improve model generalizability over diverse BMA datasets and allow for inter-institution utilization^33^. Such results also reflect the critical role of external image datasets, representing the spectrum of preanalytical variables and disease subsets, for testing any models prior to clinical deployment. Additionally, integration of cytogenetic and molecular data from patient samples could allow for cell classification models that can accurately distinguish marrow nucleated cells from samples representing various genetic subtypes of certain hematologic malignancies. Finally, as novel machine learning architectures with improved performance are developed, their utilization in automated workflows for BMA analysis will continue to improve predictive accuracy in region classification, cell detection, and cell classification models.

Overall, the automated pipeline for BMA DCCs presented here addresses and improves upon several shortcomings of manual DCCs. From WSIs, automated DCCs can utilize all identified cells, often at least two orders of magnitude more than that used for conventional 500-cell protocols, affording significantly lower imprecision in DCC values^8^, which our data suggest is at least in part due spatial cell-content heterogeneity (**Figure 5**)^9,10^. We showed that addressing both these shortcomings significantly reduces the observed variation in DCC percentages (**Figure 6**). Finally, utilizing independent models for region classification, cell detection, and cell classification trained on all available annotated data avoids inter-observer variability in counting and cytologic classification of marrow cells that ultimately lead to differences in DCC percentages^11^. Machine learning-based pipelines for automated DCCs on BMA smears present many advantages over the current gold standard of labor-intensive manual DCCs, and continued improvement with training on diverse BMA training sets representing a constellation of pathologies and morphologies, and continued updating of machine learning models, will ultimately move these pipelines into the clinical laboratory.

Supplementary information is available at Modern Pathology’s website.

## ACKNOWLEDGEMENTS

This study was supported by National Institutes of Health National Cancer Institute grants U01CA220401 and U24CA19436201.

## ETHICS APPROVAL AND CONSENT TO PARTICIPATE

This study was approved by the Institutional Review Board. The study was performed in accordance with the Declaration of Helsinki.

## AUTHOR CONTRIBUTIONS

J.E.L., L.A.D.C., and D.L.J. conceived the study; C.W.S., B.R.D, A.A., and D.L.J. identified and digitized samples; G.H.S., D.A.G., and L.A.D.C. provided computational resources and the annotation platform; J.E.L., C.W.S., B.R.D., X.Z., N.S., M.A., M.C.H., A.K., A.A., and D.L.J. created region and cell annotations; J.E.L. and L.A.D.C. developed machine learning models; J.E.L., C.W.S., B.R.D., X.Z., N.S., A.A., L.A.D.C., and D.L.J. analyzed and interpreted results; J.E.L., L.A.D.C., and D.L.J. wrote the manuscript. All authors read and approved the final paper.

## DATA AVAILABILITY STATEMENT

TensorFlow models for region detection, cell detection, and cell classification, as well as visualizations of annotations generated from the WSI BMA pipeline on testing slides, will be made available. Data sharing requests should be sent to David L. Jaye.

## SUBJECT ONTOLOGY

Bone marrow; Aspirate; Differential; Machine learning; Whole-slide imaging

