## Supplementary Information for "An Automated Pipeline for Differential Cell Counts on Whole-Slide Bone Marrow Aspirate Smears"

**SUPPLEMENTARY TABLES**

**Supplementary Table 1.** List of slides used for training and testing of models developed for the automated cell differentials pipeline, including corresponding patient diagnosis, slide cellularity, and manual cell differential values.

*Supplementary Table 1 included as Excel file.*

**Supplementary Table 2.** Replicates of manual DCCs from different evaluators on slides showing discrepancies between manual and automated blast and plasma cell percentages.

*Supplementary Table 2 included as Excel file.*

**SUPPLEMENTARY FIGURES**

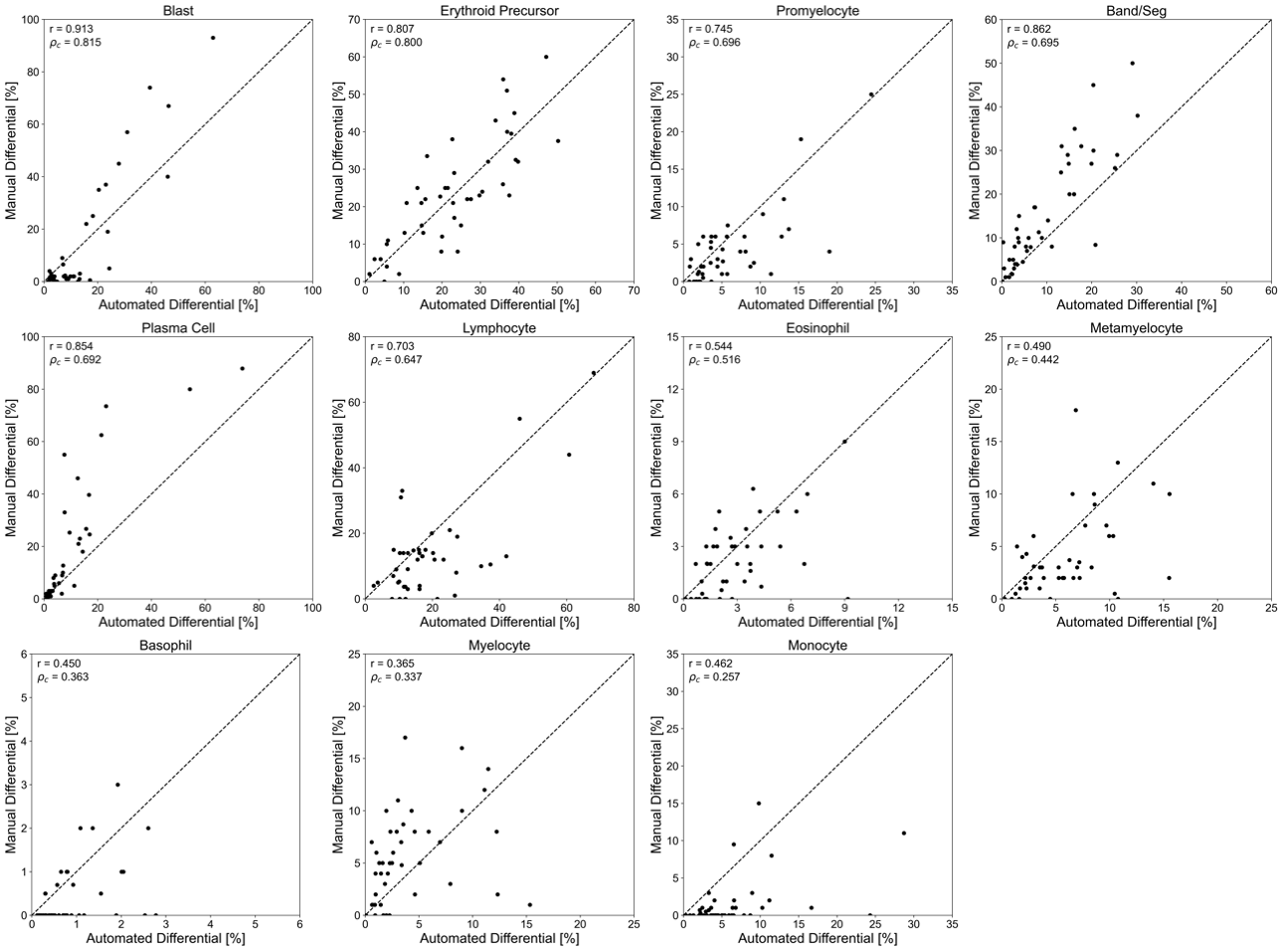

**Supplementary Figure 1.** Correlation plots comparing the percentage of each cell type obtained from the manual differential, versus the percentage obtained from the automated machine learning-based pipeline averaged across all identified viable cells in each slide. Each point represents one of 44 testing slides. Dotted line represents a 1:1 correlation between manual and automated values. r: correlation coefficient between manual and automated values. ⍴_c_: concordance correlation coefficient between manual and automated values, representing how well points are fitted by the 1:1 correlation line.

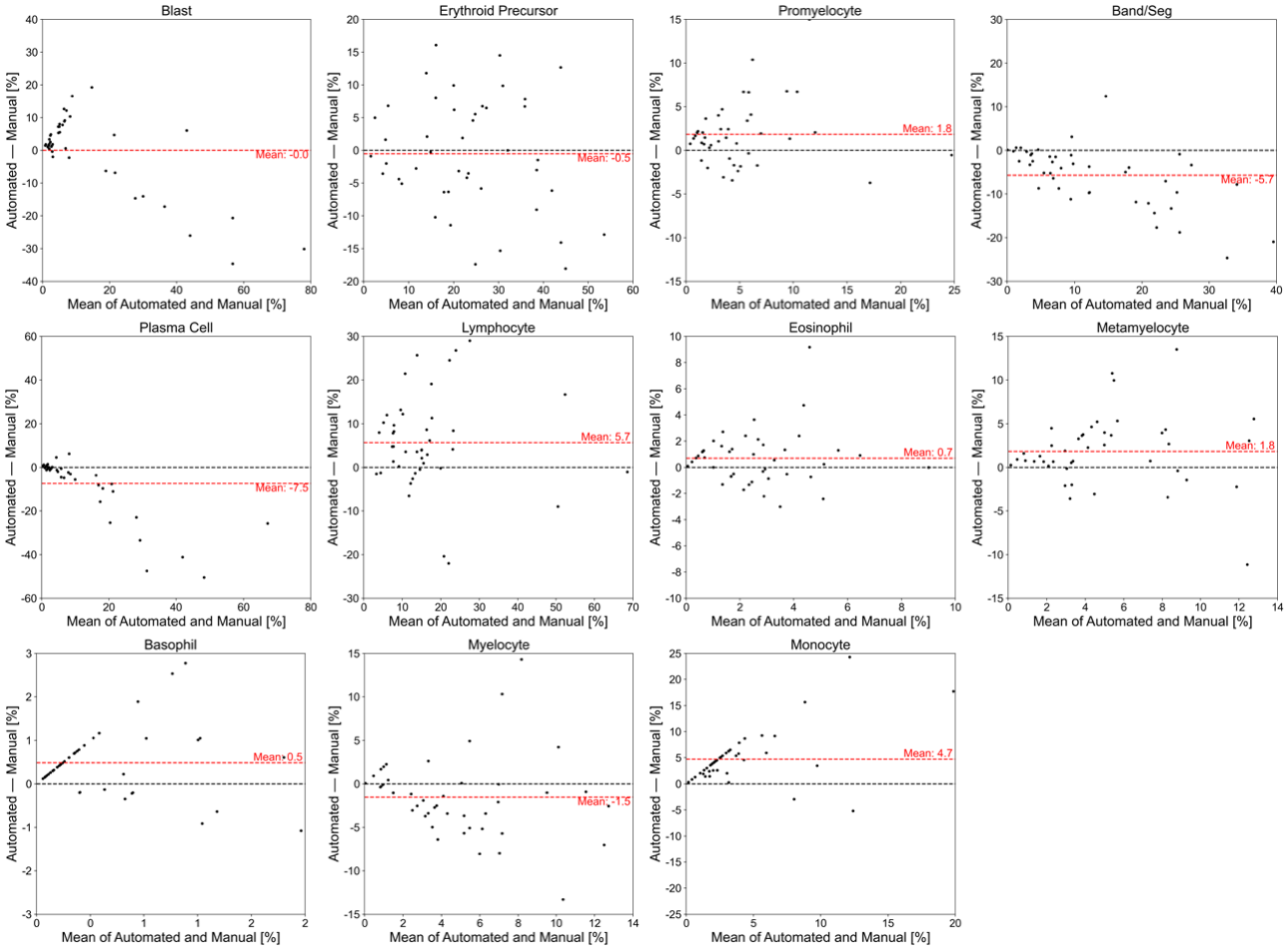

**Supplementary Figure 2.** Bland-Altman plots showing the bias between manual and automated differential values as a function of mean cell differential value.

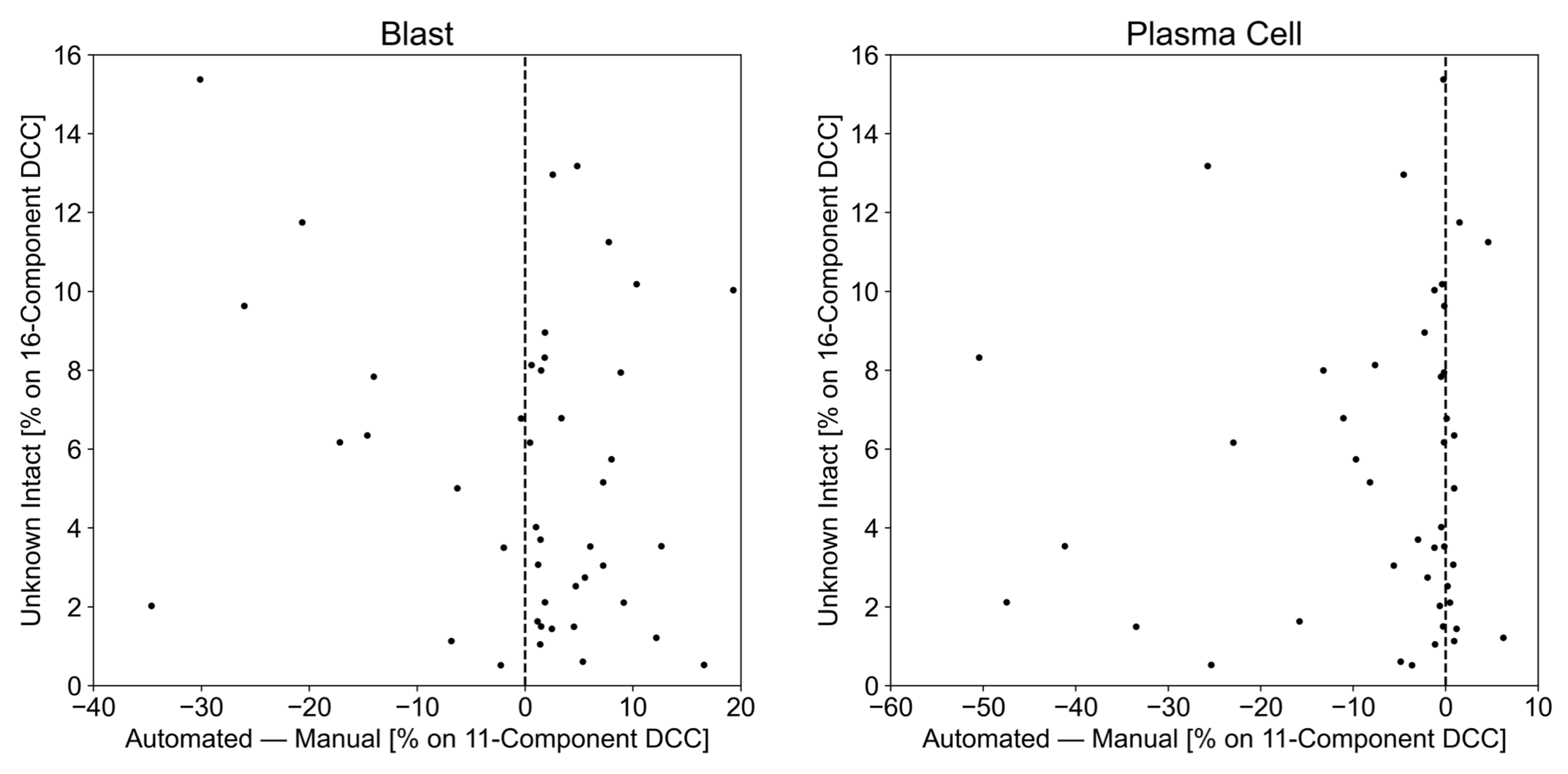

**Supplementary Figure 3.** Comparison between (x-axis) difference in automated and manual DCC values and (y-axis) calculated percentage of unknown intact cells in automated DCCs, for (left) blasts and (right) plasma cells.

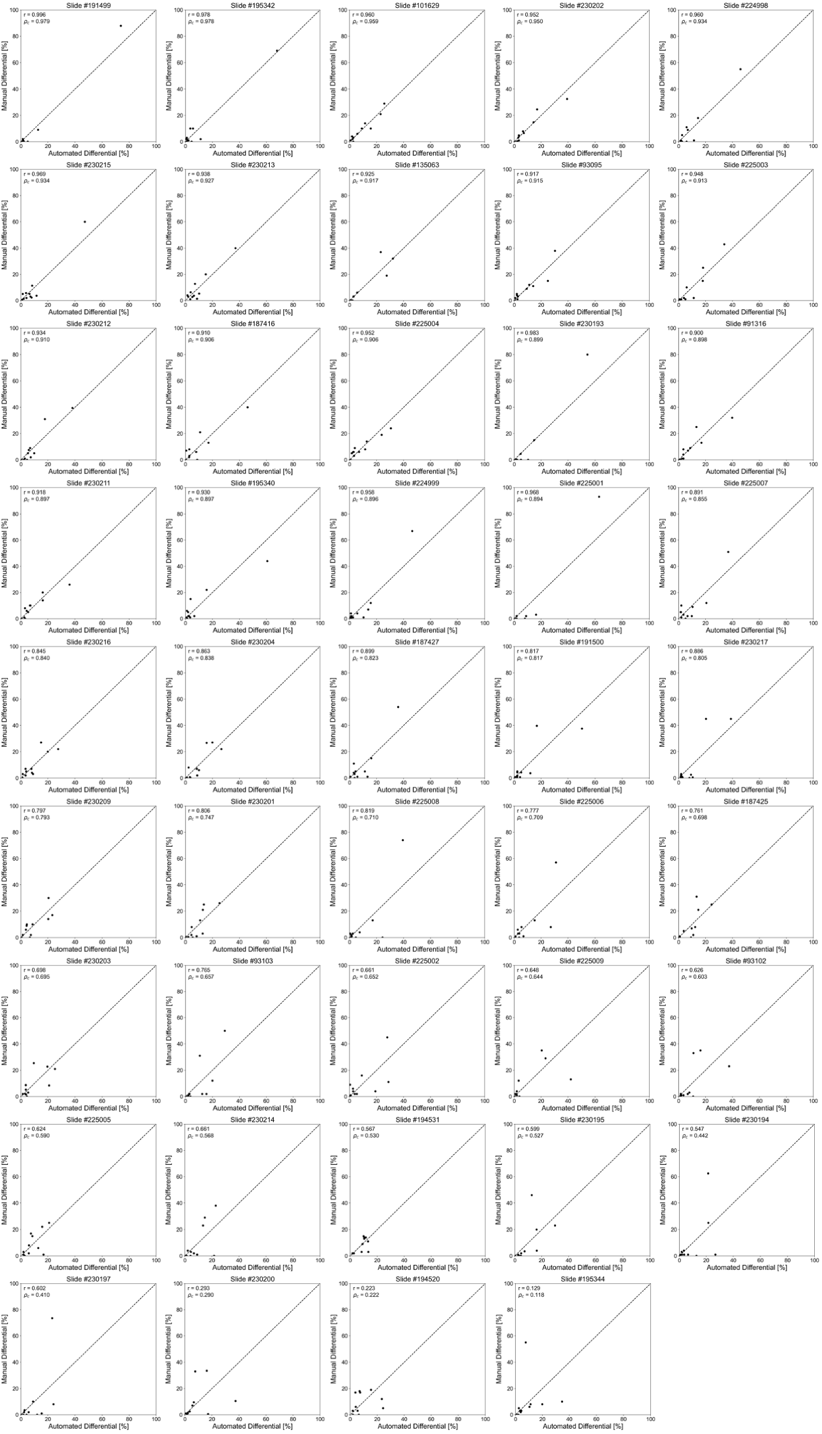

**Supplementary Figure 4.** Correlation plot comparing the percentage of 11 different cell types from the manual 11-cell differential of an individual slide, versus the percentages obtained from the automated machine learning-based pipeline. Each point represents one of 11 cell types. Vertical and horizontal error bars represent the statistically expected variability in manual and automated differential values, respectively, based on the number of cells identified from the manual and automated methods. Dotted line represents a 1:1 correlation between manual and automated values.

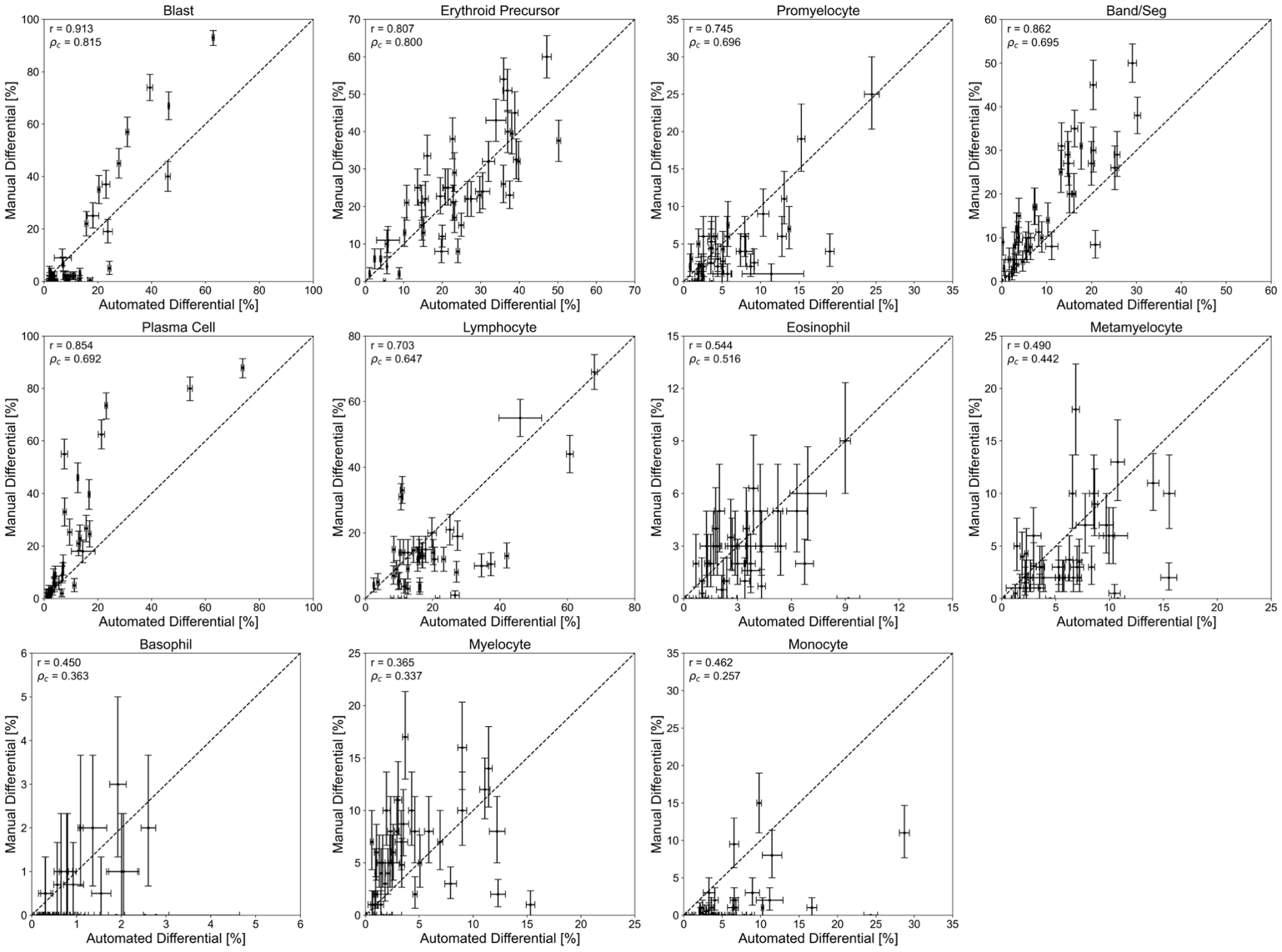

**Supplementary Figure 5.** Correlation plots cell type percentages between manual and automated DCCs, with vertical and horizontal error bars representing the statistically expected variability in manual and automated differential values, respectively, based on the number of cells identified from the manual and automated methods. Each point represents one of 44 testing slides. Dotted line represents a 1:1 correlation between manual and automated values. r: correlation coefficient between manual and automated values. ⍴_c_: concordance correlation coefficient between manual and automated values, representing how well points are fitted by the 1:1 correlation line.

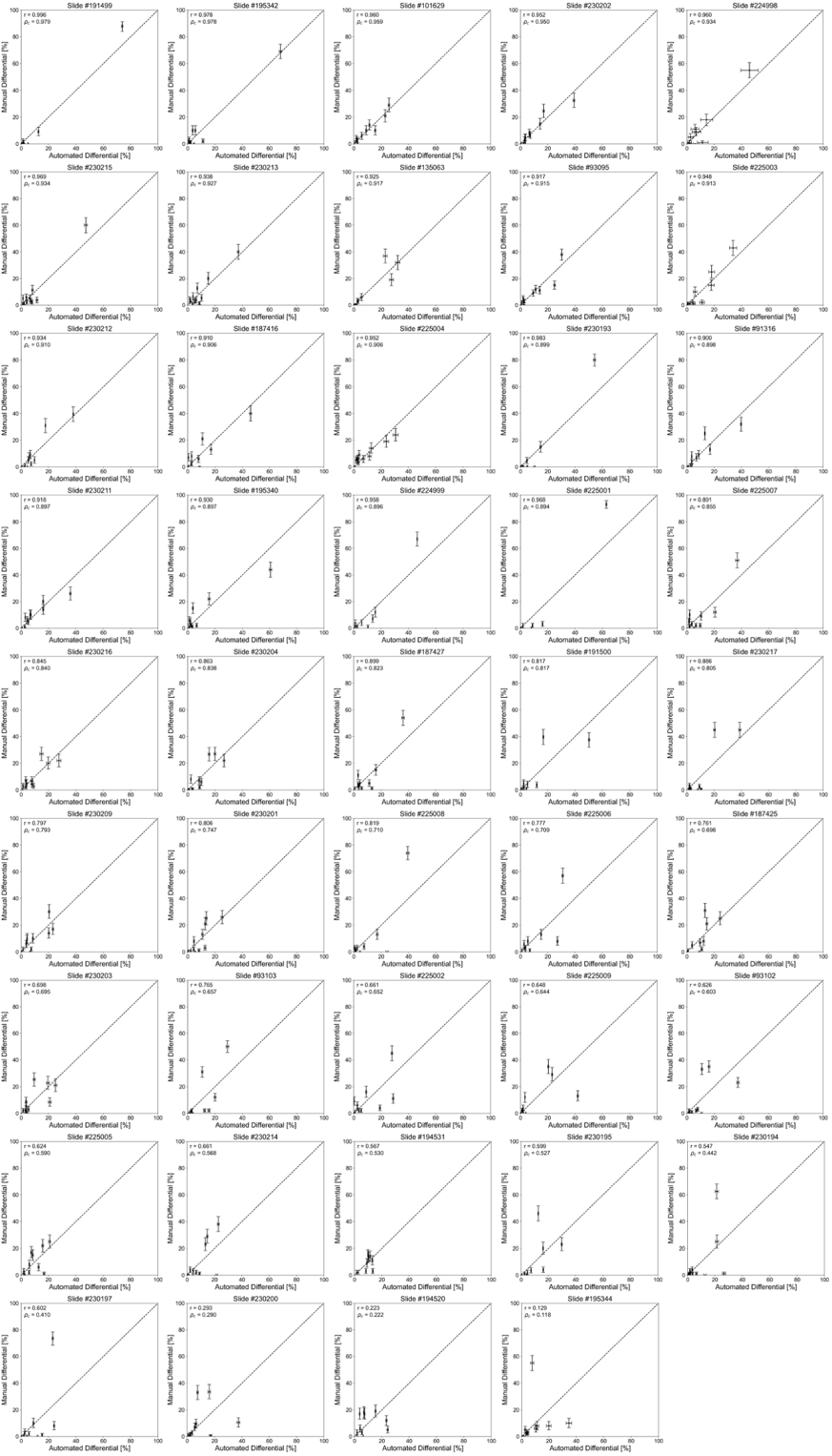

**Supplementary Figure 6.** Correlation plot comparing the percentage of 11 different cell types between manual and automated DCCs, with vertical and horizontal error bars represent the statistically expected variability in manual and automated differential values, respectively, based on the number of cells identified from the manual and automated methods. Each point represents one of 11 cell types. Dotted line represents a 1:1 correlation between manual and automated values.

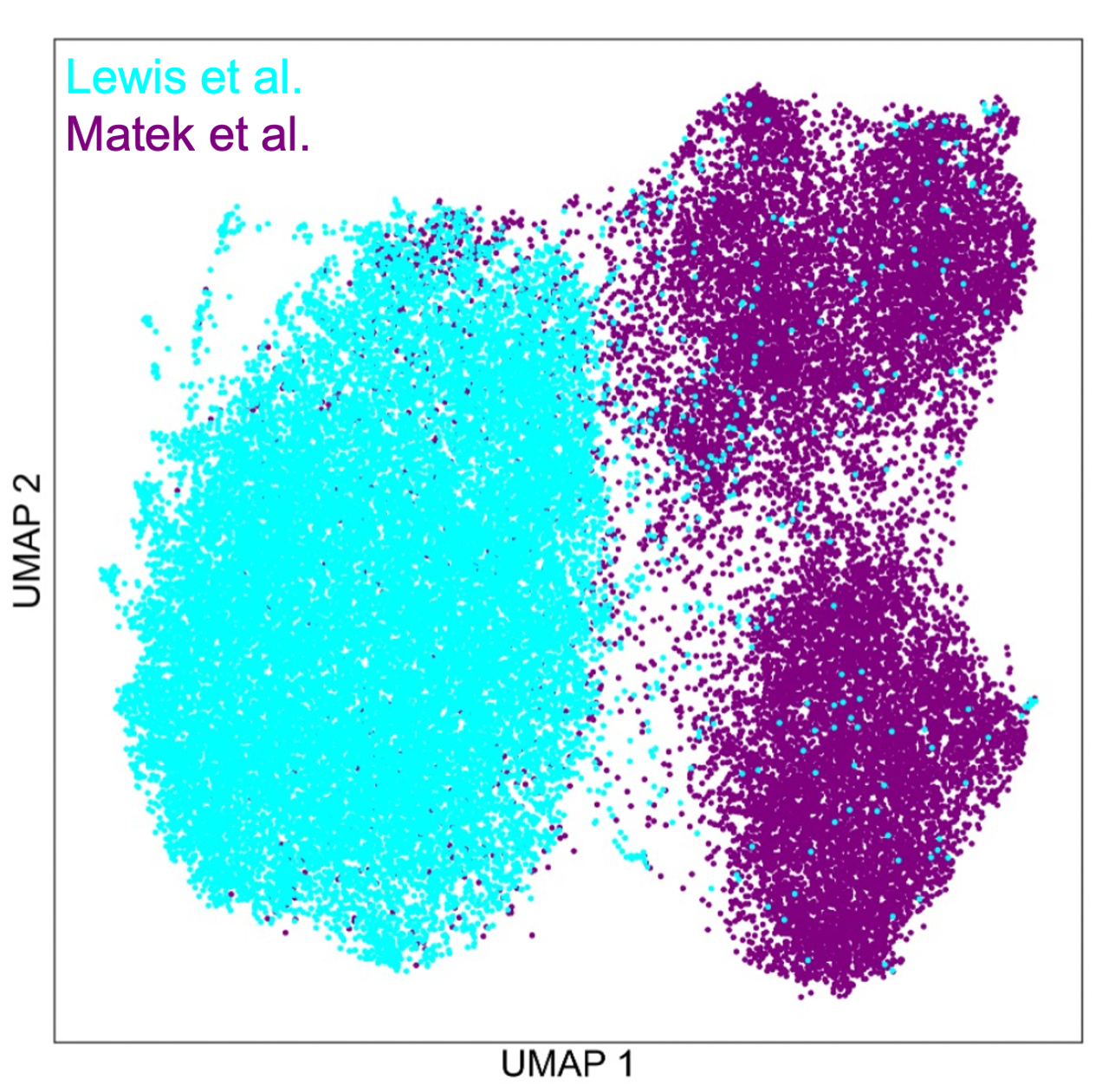

**Supplementary Figure 7.** UMAP plot comparing cell images from our cell classification training dataset (cyan) with cell images from the publicly-available cell classification training dataset presented in Matek et al. (purple).
